# High-Accuracy Multiplexed SARS-CoV-2 Antibody Assay with Avidity and Saliva Capability on a Nano-Plasmonic Platform

**DOI:** 10.1101/2020.06.16.155580

**Authors:** Tiancheng Liu, Jessica Hsiung, Su Zhao, Jessica Kost, Deepika Sreedhar, Kjerstie Olson, Douglas Keare, John Roche, Carl V. Hanson, Cynthia Press, John Boggs, Jorge P. Rodriguez-Soto, Jose G. Montoya, Meijie Tang, Hongjie Dai

**Affiliations:** Nirmidas Biotech Inc., 2458 Embarcadero Way, Palo Alto, CA 94303, USA; California Department of Public Health Viral and Rickettsia Disease Laboratory, 850 Marina Bay Parkway, Richmond, CA 94804, USA; Dr. Jack S. Remington Laboratory for Specialty Diagnostics, National Reference Center for the Study and Diagnosis of Toxoplasmosis, Palo Alto Medical Foundation, Palo Alto, CA 94301, USA; Department of Chemistry and Bio-X, Stanford University, Stanford, CA 94305, USA

**Author notes:** These authors contributed equally (Tiancheng Liu, Jessica Hsiung, Su Zhao, Jessica Kost).

## Abstract

The outbreak and rapid spread of SARS-CoV-2 virus has led to a dire global pandemic with millions of people infected and ~ 400,000 deaths thus far. Highly accurate detection of antibodies for COVID-19 is an indispensable part of the effort to combat the pandemic^1,2^. Here we developed two-plex antibody detection against SARS-CoV-2 spike proteins^3^ (the S1 subunit and receptor binding domain RBD) in human serum and saliva on a near-infrared nano-plasmonic gold (pGOLD) platform^4–8^. By testing nearly 600 serum samples, pGOLD COVID-19 assay achieved ~ 99.78 % specificity for detecting both IgG and IgM with 100 % sensitivity in sera collected > 14 days post disease symptom onset, with zero cross-reactivity to other diseases. Two-plex correlation analysis revealed higher binding of serum IgM to RBD than to S1. IgG antibody avidity toward multiple antigens were measured, shedding light on antibody maturation in COVID-19 patients and affording a powerful tool for differentiating recent from remote infections and identifying re-infection by SARS-CoV-2. Just as important, due to high analytical sensitivity, the pGOLD COVID-19 assay detected minute amounts of antibodies in human saliva, offering the first non-invasive detection of SARS-CoV-2 antibodies.

---

The severe acute respiratory syndrome coronavirus 2 (SARS-CoV-2) has rapidly spread throughout the globe within months, causing millions of infections and ~ 450,000 deaths thus far. Currently, there is no effective vaccine or drug for preventing or treating COVID-19. Diagnosis of COVID-19 has been relying on molecular detection of the SARS-CoV-2 virus by RT-PCR or isothermal nucleic acid amplification methods in a narrow time window post infection^9–12^. Antibody testing is highly complementary to molecular diagnosis and is becoming increasingly important to both the short- and long-term assessment of COVID-19 infected individuals^1,2^. Highly accurate detection of antibodies against SARS-CoV-2 is crucial to combating the pandemic by aiding diagnosis and assessing infection timing, prevalence, duration of antibody response, and potential immunity. Thus far, a variety of antibody tests have been developed using lateral flow rapid tests, enzyme-linked immunosorbent assay (ELISA) and chemiluminescence (CLIA) platforms, including ones authorized by the U.S. Food and Drug Administration (FDA) for emergency use^2,13–21^. Despite the progress, few assays differentiate antibody subtypes to glean infection timing with > 99.5 % specificity for both IgG and IgM^22,23^. High specificity is critical to avoid misinterpretations or false positives, unnecessary stress and quarantine, and to prevent controversies and wrong conclusions for surveillance or prevalence studies^23–29^. On the other hand, currently no antibody avidity test^7,30–34^ exists for COVID-19 to assess the antibody maturation and infer the timing of recent as opposed to remote infection. Antibody avidity could also aid differentiation of primary infection from secondary infection in case SARS-CoV-2 causes re-infection of recovered individuals in subsequent COVID-19 pandemic waves. Lastly, none of the SARS-CoV-2 antibody tests developed thus far offer testing of non-invasive matrices such as saliva. The non-invasive alternatives could greatly facilitate population-based mass-screening of COVID-19.

Here we developed a high accuracy, semi-quantitative assay on the nanostructured plasmonic gold (pGOLD) platform^4–8^ for detecting IgG, IgM and IgG avidity against SARS-CoV-2 spike proteins S1 subunit and RBD^1,3,35^ in human serum and saliva. The pGOLD substrate was comprised of nanoscale gold islands with abundant nanogaps, affording near-infrared (NIR) fluorescence enhancement by up to ~ 100-fold owing to plasmonic resonance and local electric field enhancements^4–8^. The greatly increased NIR signal-to-background ratio on pGOLD allowed multiplexed detection of panels of biological analytes over wide dynamic ranges. Previous antigen arrays on pGOLD simultaneously detected IgG, IgM, and IgA antibody subtypes for type 1 diabetes^5^, toxoplasmosis^8^, HDV^36^, Zika, and Dengue viral infections^7^. In particular, the pGOLD toxoplasmosis^8^ and HDV^36^ IgG antibody assays reached ~ 100 % sensitivity owing to exquisite NIR detection capabilities on the novel nanotechnology platform.

We fabricated arrays of SARS-CoV-2 spike protein S1 subunit and RBD antigen spots on a pGOLD slide in a microarray format (Fig. 1a), for capturing IgG and IgM antibodies in a sample, followed by labeling of the captured antibodies with anti-human IgG-IRDye800 and anti-human IgM-CF-647 dye (see Methods). The pGOLD biochip was then imaged by a confocal microscopy scanner in the red and NIR channels. The IgG and IgM antibodies bound to each antigen spot (Fig.1a) were analyzed through the fluorescence intensities of IRDye800 and CF-647 dye, respectively. Dilution of a pure humanized SARS-CoV-2 IgG antibody solution over 4 orders of magnitude led to signal changes by ~ 4 logs, giving an estimated antibody detection limit of ~ 1.6 ng/mL (supplementary Fig.S1,S2). A PCR-confirmed COVID-19 positive patient sample was diluted by up to 10^5^ times, and antibody signals were still well above background noise across all dilutions (Fig.S1,S2). These results suggested high analytical sensitivity and wide dynamic range of the multiplexed pGOLD assay.

**Figure 1.**
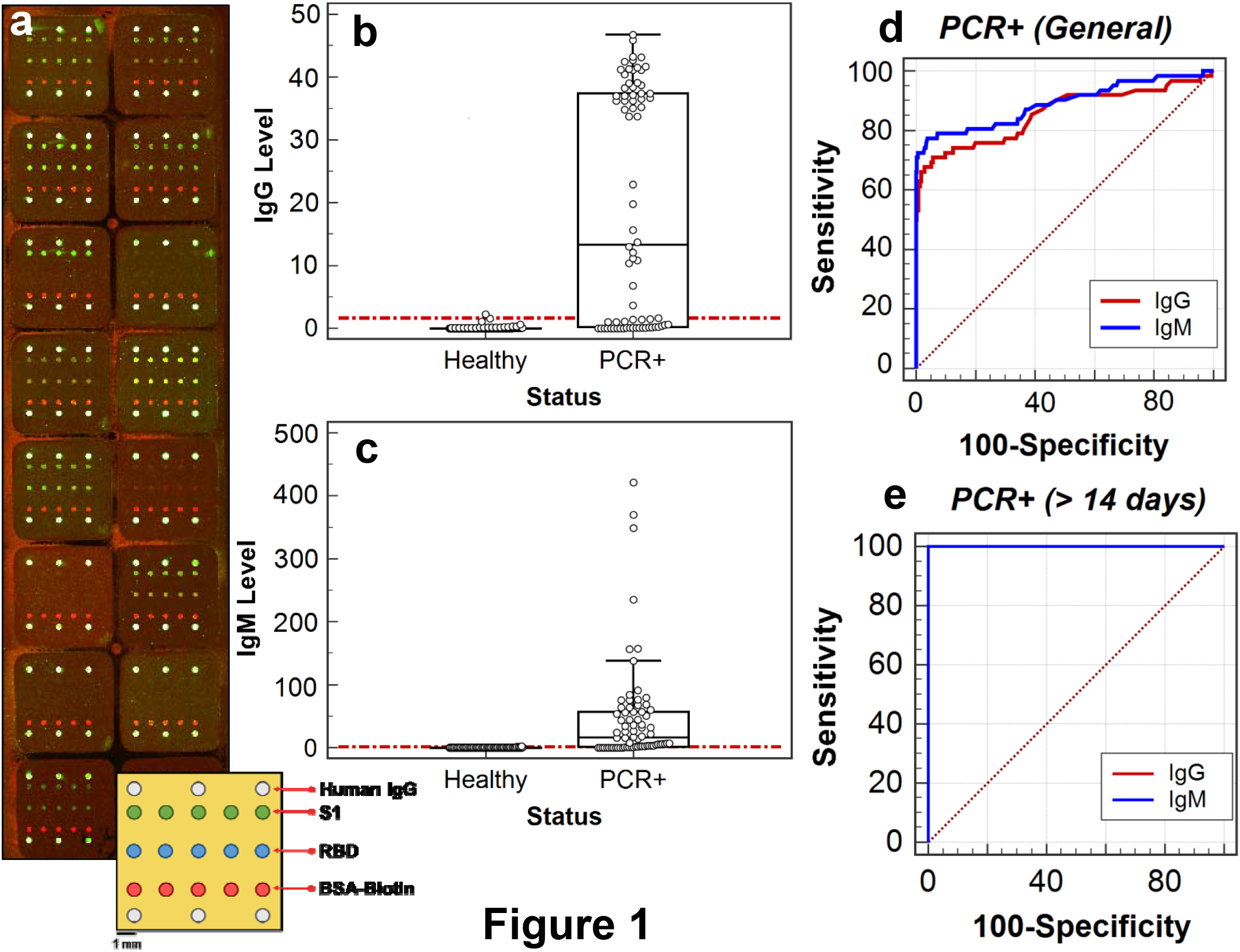
A nano-plasmonic platform for SARS-CoV-2 antibody testing. (a) An overlay of confocal fluorescence scanned images of IgG (green) and IgM (red) channels acquired after testing 16 serum samples in 16 isolated wells (square-shaped regions). Yellowish-green colored spots correspond to the presence of both IgG and IgM in the sample. The lower right schematic drawing shows the printing layout of S1 (in green) and RBD (in blue) antigens and human IgG control spots (in white) in each well. The BSA-biotin spots (in red) are always labeled by a streptavidin dye in the IgM fluorescence channel to serve as an intrawell signal normalizer. (b) Box plots of IgG levels detected in PCR-negative COVID-19 or presumptive negative (‘Healthy’) and PCR-positive (‘PCR+’) COVID-19 samples with the cutoff indicated as a dashed red line. (c) The same as (b) except for IgM. (d) ROC curve for pGOLD SARS-CoV-2 IgG/IgM assay based on 384 negative and 62 PCR-positive COVID-19 serum, which was used to establish IgG and IgM cutoffs. (e) ROC curve for pGOLD SARS-CoV-2 IgG/IgM assay based on 384 negative and PCR-positive COVID-19 serum samples collected 15-45 days post symptom onset.

We detected SARS-CoV-2 antibodies on the pGOLD assay in human serum samples, first focusing on antibodies against the S1 antigen (Fig.1). To determine specificity, we tested a total of 384 negative and presumptive negative samples including 33 from PCR-confirmed COVID-19 negative individuals, 311 pre-pandemic samples collected in 2017-2019, and 40 healthy control samples acquired prior to the COVID-19 outbreak. We also obtained a set of sera from 62 PCR-confirmed COVID-19 patients (but without information given regarding the number of days between disease symptom onset to sample collection). The ROC (receiver operating characteristics) curve analysis was performed based on the pGOLD assay results for the 384 negative and 62 positive samples (Fig.1d). The cutoff values were determined under the criteria of > 99.5 % specificity while maximizing the sensitivity for detecting both IgG and IgM in the sera of COVID-19 patients (Fig.1b,1c). Under this condition, only one serum sample from the 384 presumptive negative set was found to be false positive.

To further establish pGOLD assay specificity and potential cross-reactivity, we tested 70 pre-pandemic samples collected from patients with various diseases, including common colds/other coronaviruses, influenza, autoimmune disease, HBV, HCV, and HIV (Table S4). All of the samples were found negative in IgG and IgM against SARS-CoV-2, suggesting near zero cross-reactivity of the pGOLD assay (Fig.2c,2d). All together with a total of 454 pre-pandemic presumptive and PCR-confirmed negative samples (Fig.2c,2d, Table S1), only one sample was false positive in IgG and IgM, resulting in an overall specificity of 99.78% for both antibody isotypes.

**Figure 2.**
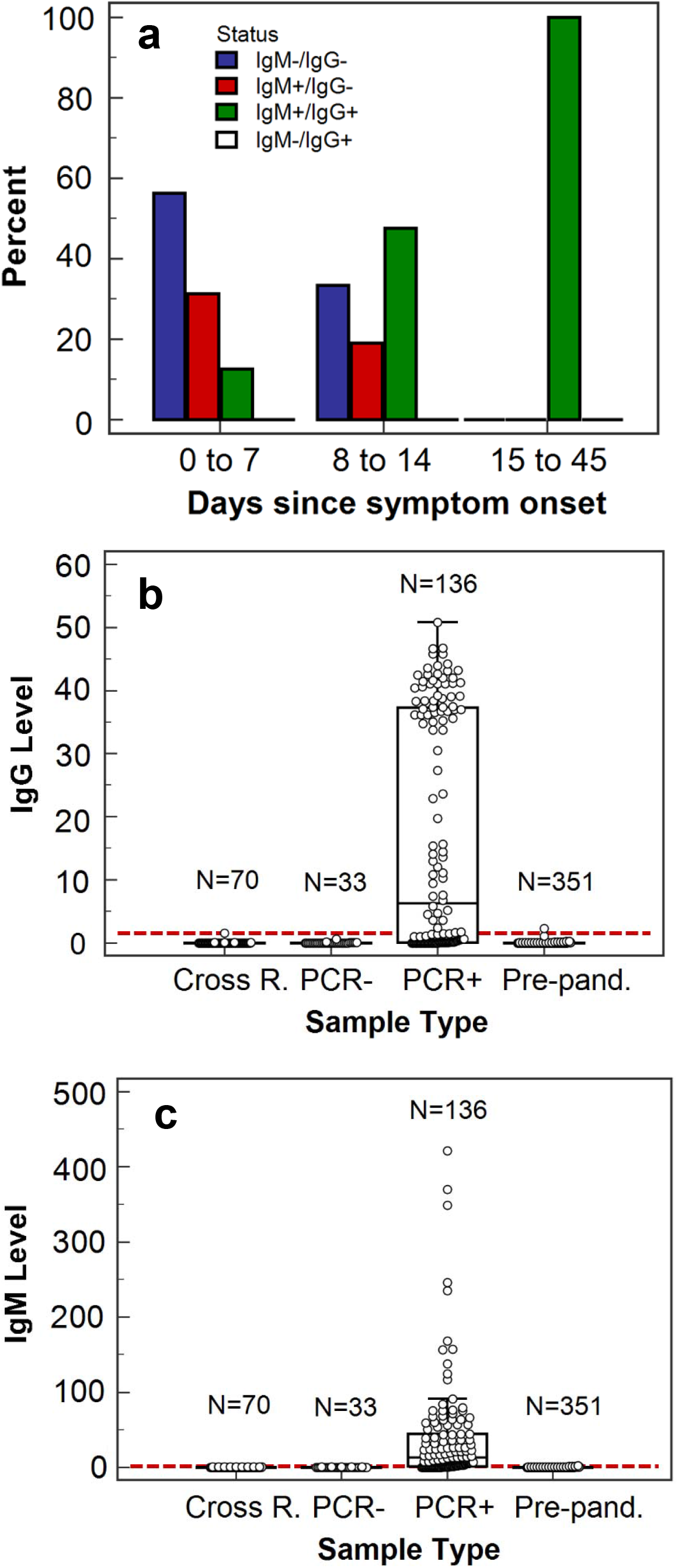
Highly sensitive and specific SARS-CoV-2 antibody test. (a) Percentages of samples with IgG/IgM antibody status combinations according to days from symptom onset to sample collection date in a range from 0-7, 8-14, and 15-45 days. (b) Box plots of IgG levels detected in four groups of serum samples indicated on the x-axis with the cutoff displayed as a dashed red line. ‘PCR+’ denotes serum samples from patients who tested positive by PCR for COVID-19 and ‘PCR-’ denotes those who tested negative. ‘Pre-pand.’ corresponds to pre-pandemic collected samples. ‘Cross R.’ corresponds to samples from patients with other diseases for cross-reactivity evaluation. (c) The same as (b) except for IgM.

For the 62 PCR-confirmed COVID-19 patient samples collected without days since symptom onset data (Fig.1), the sensitivity of the pGOLD assay was 51.61 % for IgG and 70.97 % for IgM (Fig.1d), suggesting a substantial fraction of the samples was collected in the early stage (≤ 14 days) of SARS-CoV-2 infection prior to the development of antibodies^2,13–21^. To investigate the immune response and sensitivity of pGOLD SARS-CoV-2 antibody assay at various times post infection, we measured an independent set of sera from 70 PCR-confirmed COVID-19 patients collected between 0-45 days post symptom onset. Assessment of antibody status in the second set of samples was performed using the cutoffs generated with the first sample set. Based on the days of symptom onset, PCR-positive patients were divided into three groups: I (0-7 days), II (8-14 days) and III (>14 days). We found that the positive rate for IgM in each group (I, II, III) was 43.75 %, 66.67 %, and 100 %, respectively (Table S2). The positive rate for IgG in each group (I, II, III) was 12.5 %, 47.62 %, and 100 %, respectively (Table S2). The results indicate a high positivity rate of IgM over IgG initially (Group I), and both IgG and IgM antibodies detected in all patients at a later stage post infection (Group III).

In Group I (0-7 days post infection), about 56.25 % of patients were negative for both IgM and IgG, 31.25 % of patients developed IgM but not IgG, and 12.5 % of patients developed IgM and IgG, clearly showing the presence of IgM preceding IgG as the initial immune response against COVID-19 infection. In Group II, 47.62 % of patients developed both IgM and IgG compared to 19.05 % of patients who were positive for IgM and negative for IgG within the same onset group. By > 14 days post infection, 100% patients developed both IgG and IgM against SARS-CoV-2 (Fig.2a). Combined, the antibody positive rate for ≥ 6 days, ≥ 10 days and > 14 (15-45) days were 75% for IgG and 86.67% for IgM, 87.76% for IgG and 93.87% for IgM, and 100% for IgG and 100% for IgM, respectively (Table S3).

The pGOLD SARS-CoV-2 antibody assay clearly observed the classical serological immune response behavior of an earlier appearance of IgM before IgG post infection in the 0-14 days acute phase. This differed from a recent serological-response study by ELISA that found the IgM positive rate below IgG at all times^20^. The low positive rate of IgM by ELISA could be due to insufficient analytical sensitivity, causing delays in the detection of IgM levels above background noise in PCR-confirmed COVID-19 patient samples^37,38^. The pGOLD assay has near 100% sensitivity (for COVID-19 patient sera collected > 14 days post infection) and 99.78 % specificity in detecting both IgG and IgM subtypes (based on ~ 550 presumptive negative and positive samples, Fig.1e,2b,2c). Most of the FDA-authorized instrument-based laboratory diagnostic tests are specific and sensitive but do not differentiate antibody subtypes, and none of the lateral flow rapid tests can match the sensitivity of laboratory tests (EUA authorized serology test performance: https://www.fda.gov/medical-devices/emergency-situations-medical-devices/eua-authorized-serology-test-performance).

In addition to exceptional analytical sensitivity, the pGOLD assay was capable of testing IgM and IgG antibodies in the same patient sample against multiple SARS-CoV-2 antigens simultaneously (Fig.1a), a unique feature among existing COVID-19 assays. To investigate antibody responses to specific regions of the SARS-CoV-2 spike protein, we measured and analyzed antibodies against RBD in the S1 subunit. We obtained a maximum specificity of 99.78% for IgG (1/454 false positive, the same anti-S1 IgG false positive) and 99.78% for IgM (1/454 false positive, a different sample from the anti-S1 IgM false positive), and sensitivity of 100% for both IgG and IgM in PCR-confirmed COVID-19 patient samples at > 14 days post disease symptom onset, suggesting that the pGOLD assay of antibodies against RBD was also highly specific and sensitive (Table S6,S7).

Correlation analysis of antibodies in COVID-19 sera against S1 and RBD deviated from linear relations, with IgG levels against the two antigens scattered around the slope = 1 line (Fig.3a). The IgM levels against S1 and RBD were scattered with a fit-line slope well below 1, suggesting substantially higher IgM binding to RBD than to S1 (Fig.3b). The RBD in the S1 subunit in the SARS-CoV-2 spike protein was responsible for binding to the ACE2 receptor on host cells to initiate infection^3,39^, and was generally recognized as an important target for both neutralization antibody treatment and specific SARS-CoV-2 detection. Our results showed that the S1 and RBD antigens were both highly specific and sensitive SARS-CoV-2 antibody targets, while complementing each other in sensitivity to a discernable degree (Table S5). To exploit the two-plex capability, we were able to combine antibodies against S1 and RBD to increase the pGOLD assay sensitivity for the ≤ 14-day sample group for IgG and IgM, from 32.43 % against S1 only to 37.84 % combined and 54.05 % against RBD only to 56.76% combined, respectively.

**Figure 3.**
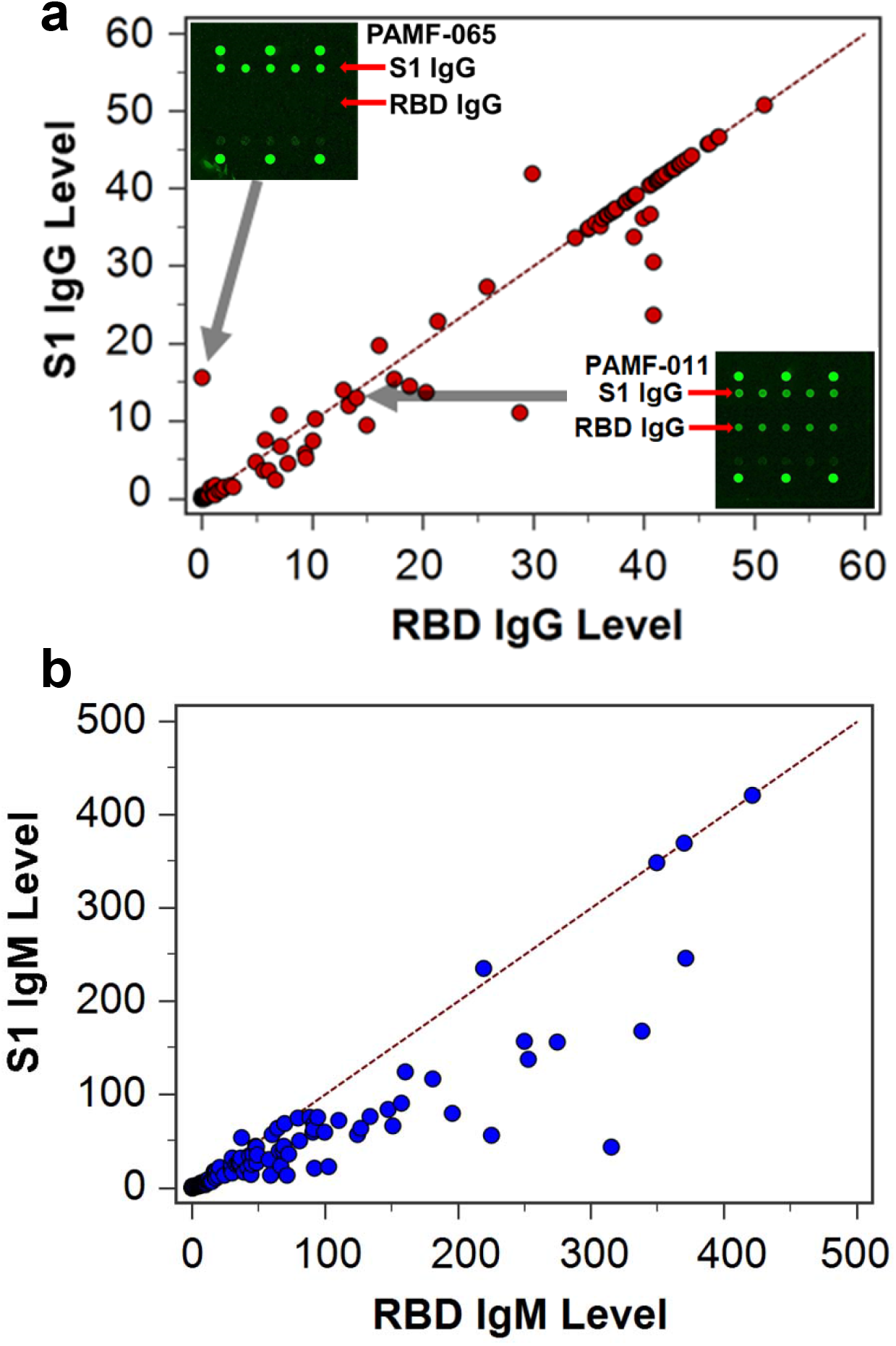
Correlation of antibodies against two SARS-CoV-2 antigens. (a) Correlation plot of anti-S1 IgG level (y-axis) and anti-RBD IgG level (x-axis) measured in PCR-confirmed COVID-19 patient sera. The dashed line was drawn to have a slope of 1. The upper left inset shows the scanned image of the IgG-only channel in a patient serum labeled as PAMF-065, which displayed high signal on the S1 antigen but not on the RBD antigen. The lower right inset shows the scanned image of IgG levels of a sample labeled as PAMF-011, displaying about equal IgG signals against S1 and RBD. (b) Correlation plot of anti-S1 IgM level (y-axis) and anti-RBD IgM level (x-axis) measured in COVID-19 patient sera. The dashed line was drawn to have a slope of 1.

Intriguingly, an extreme case was that a COVID-19 patient serum (labeled PAMF-065) showed SARS-CoV-2 IgG only binding to S1 and not to RBD (Fig.3a). A recent study identified distinct groups of phage-display derived antibodies against SARS-CoV-2 that bind preferentially to RBD or S1^40^. Our results could reflect a similar phenomenon in COVID-19 human serum, but requires further understanding of immune responses to SARS-CoV-2 and the antibody-antigen interactions at the molecular level^41^.

A SARS-CoV-2 IgG avidity assay was developed on pGOLD to investigate antibody-antigen binding affinity and stability in denaturing conditions^7,30–34^. Antibodies developed shortly after a primary infection exhibit low avidity and bind weakly to the antigen. Overtime avidity towards antigens can increase as antibodies ‘mature’ through colonial expansion, hypermutation and affinity selection in the germinal center^42^. IgG avidity has been previously used to aid differentiation of recent from past infection and distinguish primary from secondary infection^7,30–33^. This could be particularly important if COVID-19 returns in subsequent waves and in upcoming influenza seasons. In our pGOLD SARS-CoV-2 IgG avidity test, a denaturing 6 M urea-treatment step was introduced to remove weakly bound antibodies on the antigen, leaving only the antibodies with a strong affinity for the antigens on pGOLD to be detected.

We found that IgG against S1 and RBD in all COVID-19 patient sera except for one (PAMF-065) showed low avidity between 0 and 0.3 (Fig.4a, Fig.4b lower image, Table S8), consistent with recent infections since all of our IgG-positive COVID-19 samples (49/70 PCR-positive tested for avidity) were collected within 6-45 days post infection. A slight trend of higher anti-S1 IgG avidity vs. the number of days of post symptom onset was discerned (Fig.4a). Also noticeable was a lower average anti-RBD IgG avidity than anti-S1 IgG avidity (Fig.4a). However, testing of a large number of samples from recovered COVID-19 patients over long periods of time (> 6 months to 2 years) is needed to glean a clearer picture. In addition to assessing recent/remote infection and primary/secondary infection, multiplexed measurements of avidity towards a panel of antigens will be useful to understanding immune responses to SARS-CoV-2 and how antibodies mature post infection, with implications to immunity and convalescent plasma based antibody therapy^43^.

**Figure 4.**
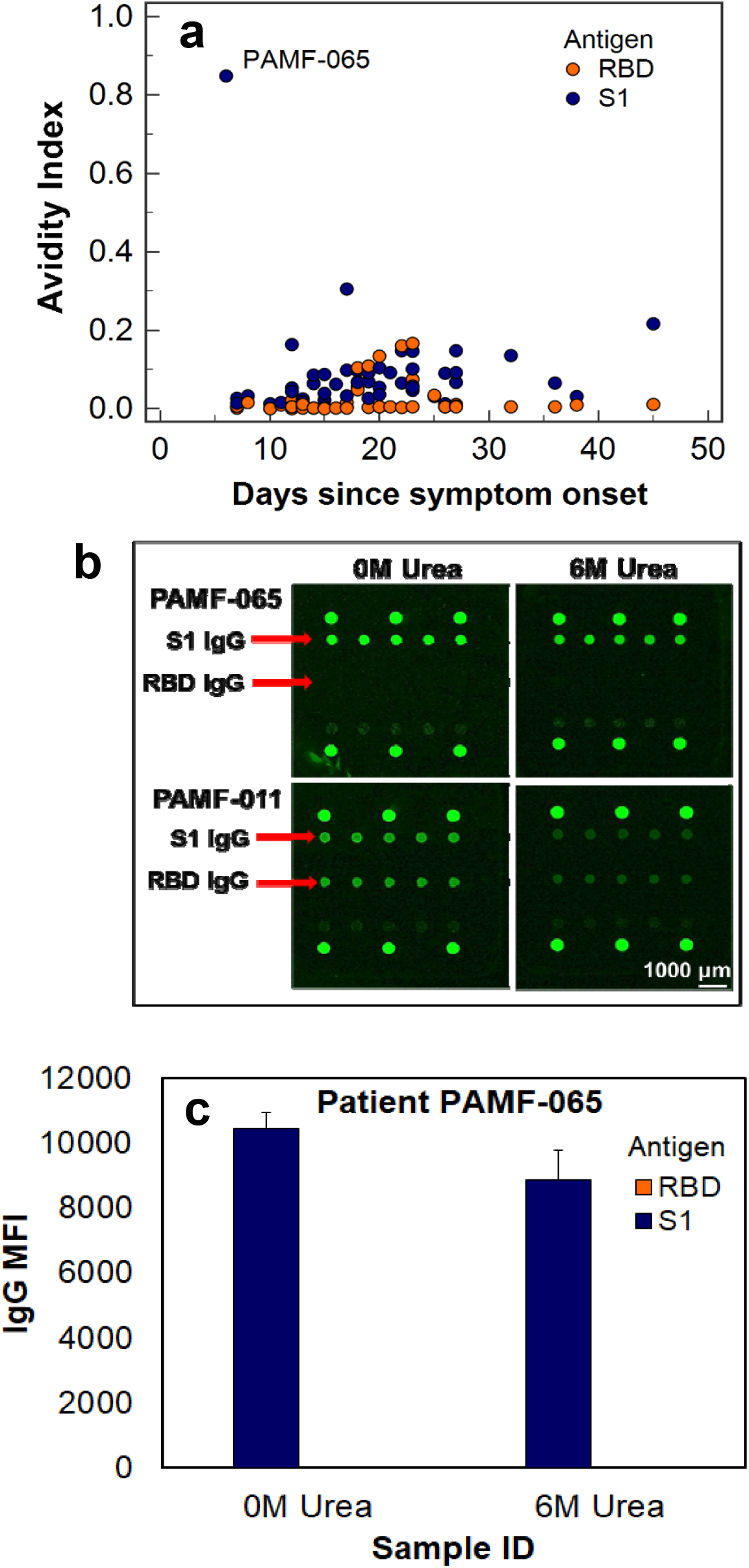
Antibody avidity against SARS-CoV-2 antigens. (a) Avidity of anti-S1 IgG and anti-RBD IgG measured in IgG-positive, PCR-confirmed COVID-19 patient sera collected 6-45 days post symptom onset. The serum of PAMF-065 showed unusually high avidity for anti-S1 IgG while being negative for anti-RBD IgG. (b) Upper panel: Fluorescence images of IgG-only channel showing PAMF-065 serum sample with high anti-S1 IgG level with and without urea treatment, hence high avidity. It showed negligible anti-RBD IgG. Lower panel: Fluorescence images showing another patient serum tested, PAMF-011, with much reduced anti-S1 IgG level after urea treatment, indicating low avidity. Low avidity was observed for all samples except PAMF-065. (c) Anti-S1 IgG median fluorescence intensity (MFI) signals of the PAMF-065 sample with and without urea treatment. The error bars indicate one standard deviation away from the mean.

Surprisingly, the sample PAMF-065 (with antibody binding only to S1 and not to RBD in Fig.3a,4b) collected from a PCR-confirmed COVID-19 patient showed a strong anomaly in anti-S1 IgG avidity with a high value of ~ 0.8 (Fig.4b upper image, Fig.4c), typically interpreted as infection > 6 months ago. The serum sample used for pGOLD antibody assay was collected from the patient only ~ 6 days post COVID-19 symptom onset, yet showed a high anti-S1 IgG level at ~ 10 times of cutoff with only a low positive IgM against S1. These characteristics were consistent with secondary infection as found in the flavivirus field^7,44^. Secondary flaviviral infection led to rapid IgG increase within days post symptom onset accompanied by high IgG avidity and low IgM levels^7,44^. The patient was a 73 years old woman tested SARS-CoV-2 positive by PCR at 6 days (same day as serum sample was obtained) post the onset of symptoms (fever, lymphopenia). Prior to the diagnosis the patient had likely been exposed to SARS-CoV-2 for several weeks from her mother who died of COVID-19. Notably that the patient did not develop pneumonia or get much sicker despite her relatively advanced age, suggesting a degree of immunity. We tentatively assign the PAMF-65 patient to re-infection based on the high IgG against SARS-CoV-2 S1 at 6 days post symptom onset, low positive IgM, and unusually high IgG avidity. It was possible that the patient was previously exposed to a closely related infection including SARS-CoV-1 with antibodies cross-react with SARS-CoV-2^45^. This case was intriguing and underscored the usefulness and importance of antibody avidity testing for COVID-19.

Lastly, we exploited the high analytical sensitivity of the nano-plasmonic gold platform for detecting antibodies in human saliva against SARS-CoV-2. It is well known that antibody concentration in human saliva is orders of magnitude lower than in blood or serum, demanding assay platforms with exquisite analytical sensitivity and capable of detecting ultra-high signal over background noise^8^. We tested the saliva samples of 4 fully recovered, PCR-positive COVID-19 patients and 11 healthy non-infected individuals on pGOLD (Fig.5a). Although background fluorescence signals were observed for some saliva samples, likely due to autofluorescence of molecules in the saliva, background-subtracted IgG signals on S1 and RBD antigens allowed clear differentiation of positive COVID-19 recovered patient saliva from healthy controls (Fig.5b). This promising result suggested the first multiplexed saliva-based antibody test for SARS-CoV-2, which could greatly facilitate population-based mass screening of COVID-19. Note that a COVID-19 patient serum diluted by 10^4^ times was included in the assay (labeled ‘REF’ in Fig.5) and showed similar IgG level as in the COVID-19 positive saliva samples.

**Figure 5.**
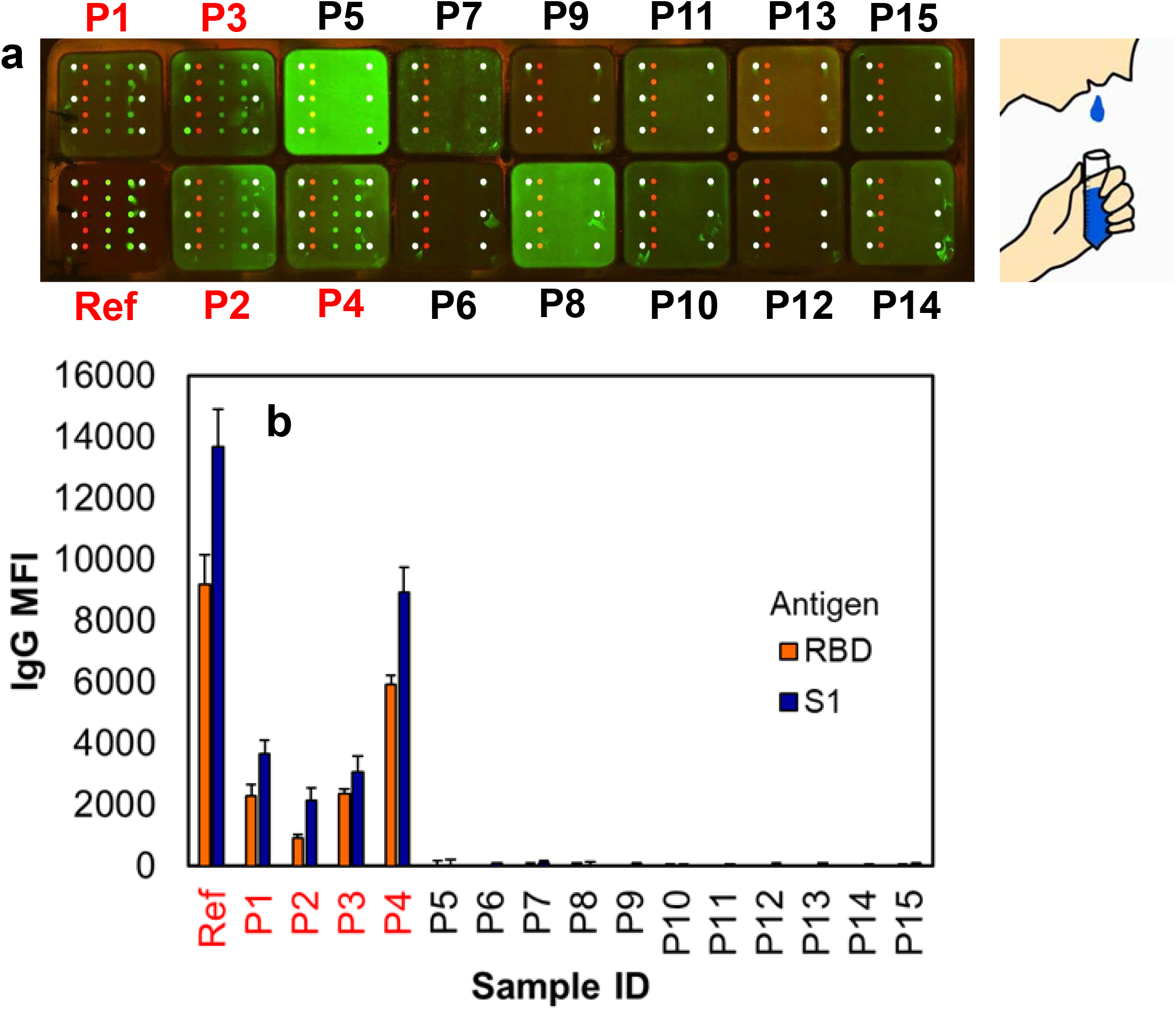
Detection of SARS-CoV-2 antibodies in human saliva. (a) A confocal fluorescence image of IgG signals in the saliva of 4 recovered COVID-19 patients (denoted as P1-P4) and 11 healthy controls (denoted as P5-P15) and a 10^4^ times diluted serum of a PCR-confirmed COVID-19 patient as a reference (denoted as ‘Ref’). Saliva was collected by a simple spitting method as shown in the schematic. (b) Median fluorescence intensity (MFI) signals of anti-S1 and anti-RBD IgG measured in the saliva samples and PCR-positive COVID-19 serum reference with background signals subtracted. The error bars indicate one standard deviation away from the mean.

With ~ 100% sensitivity two weeks post infection and 99.78% specificity based on > 450 negative samples, the pGOLD COVID-19 IgG/IgM assay is promising for population-based mass screening, and sero-surveillance and prevalence studies. The clear trend of IgM detection prior to IgG could be used to aid COVID-19 diagnosis starting from the early stage. A highly sensitive and specific IgM test could facilitate diagnosis of re-infection or secondary infection in the acute phase in a future return of SARS-CoV-2, much like the utility of Zika IgM testing in flaviviral endemic regions^46^. Multiplexed IgG avidity measurements against multiple virial antigens could facilitate understanding of the immune responses and antibody maturation, and aid the differentiation of primary from secondary infection, and reveal the infection timing^47^. A unique feature is the high capability of SARS-CoV-2 antibody detection in human saliva samples on the novel nanotechnology based pGOLD platform, which can enable non-invasive home sample collection for mass-screening of COVID-19.

## Methods

### Biological samples and materials

74 PCR-confirmed COVID-19 patient serum samples were provided by the California Department of Public Health (CDPH) and the Dr. Jack S. Remington Laboratory for Specialty Diagnostics (JSRLSD) at the Palo Alto Medical Foundation. These samples were provided with information on the number of days between sample collection and disease symptom onset, excluding for 4. Another set of PCR-confirmed COVID-19 sera (with no known information on the number of days between disease onset to sample collection) were obtained from Baptist Health South Florida and Loma Linda Medical Center. 33 PCR-negative samples were provided by CDPH and Loma Linda Medical Center. 311 pre-pandemic serum samples collected in 2017-2019 were from the JSRLSD lab. 40 healthy control samples were acquired from the Arizona State University Health Services for projects before the SARS-CoV-2 outbreak. 70 samples from patients with various diseases for cross-reactivity checking were provided by Loma Linda Medical Center, CDPH, Valley Medical Center in San Jose, the JSRLSD lab, or purchased commercially. Saliva samples were collected through a simple spitting method into a plastic tube (Fig 5a) from healthy donors and fully recovered COVID-19 patients who tested positive by PCR over a month before collection. Saliva was diluted two times and centrifuged to remove any aggregates, and the supernatant was tested on the pGOLD assay.

We conjugated IRDye800 CW NHS ester (LI-COR Biosciences) and CF647 NHS ester (Millipore Sigma, SCJ4600048) to anti-human IgG and anti-human IgM, respectively. The IRDye800-labeled anti-human IgG and CF-647-labeled anti-human IgM were used for two-color simultaneous detection of IgG and IgM against S1 and RBD antigens on pGOLD.

### Multiplexed SARS-CoV-2 microarray printing on pGOLD slides

Each pGOLD slide (Nirmidas Biotech Inc.) was printed with two SARS-CoV-2 antigens, namely the spike protein S1 subunit (S1) and S1 containing the receptor binding domain (RBD), using a GeSiM Nano-Plotter 2.1 at the following concentrations: 60 μg/mL for S1 (40591-V08H, Sino Biological Inc.) and 25 μg/mL for RBD (40592-V08H, Sino Biological Inc.). On the same biochip, 7.5 μg/mL human IgG and 50 μg/mL BSA-biotin (Thermo Fisher Scientific) were also printed to serve as a printing control and “intra-well signal normalizer”, respectively. The antigens were printed in quintuplicate for capturing SARS-CoV-2 antibodies in either serum, plasma, whole blood, or saliva (the current work focused on serum and saliva). Identical microarrays were printed on 16 isolated wells in each pGOLD biochip with a total of 4 biochips resembling a 64-well plate using the FAST frame incubation chamber (Millipore Sigma). The SARS-CoV-2 antigen-printed biochips were vacuum sealed and stored at −80 °C until use.

### Multiplexed pGOLD SARS-CoV-2 IgG/IgM assay procedure

Serum samples were heated at 56 °C for 30 minutes to deactivate and reduce potential risk from any residual virus^48–50^. The heat-deactivated serum samples were immediately used or stored at −80 °C for later use. The pGOLD antibody assays were performed in a 16-well format in the following steps: **1)** Blocking: All wells were blocked with a blocking buffer for 30 minutes at room temperature, **2)** Sample incubation: each well was then incubated with 100 μL of diluted patient serum (200X diluted in a dilution buffer) or saliva (2X dilution) for 60 minutes at room temperature. A positive control (diluted patient serum) and blank control (dilution buffer only) were also included in each biochip. **3)** Secondary antibody incubation: each well was subsequently incubated with a mixture of 4 nM IRDye800-labeled anti-human IgG secondary antibody, 4 nM CF-647-labeled anti-human IgM secondary antibody, and 6 nM CF-647-labeled streptavidin for 30 minutes at room temperature (Fig. 1a). Note that each well was washed three times with PBST (PBS with 0.05 % Tween 20) between steps. The CF-647-labeled streptavidin in the detection step binds to BSA-biotin spots in each well on the pGOLD biochip, and the signal was used as an “intrawell signal normalizer”. That is, the IgG and IgM signals were divided by the intrawell normalizing signal to obtain a ratio index for IgG and IgM of each sample. Intrawell normalization was designed to minimize the effect of slight differences in pGOLD film uniformity across each biochip which may affect fluorescence enhancement.

### pGOLD SARS-CoV-2 IgG avidity assay

The pGOLD SARS-CoV-2 IgG avidity in a serum sample was measured by detecting captured IgG for the sample with and without urea treatment side-by-side in two neighboring pGOLD wells. In each well, 0.5 μL of the same serum sample was diluted 200 times. In one well, a regular IgG assay was performed, whereas in the neighboring well the same IgG assay was performed except that a 10-minute treatment with 6 M urea was added following the sample incubation step. Such treatment by a denaturing agent like urea could detach the IgG from the antigen spot if the IgG avidity is low. At the end of the assay, the IgG signal of the urea-treated sample was divided by the IgG signal of the regularly assayed sample, giving an avidity index value.

### Data analysis

After the assay procedures, a dual-channel MidaScan microarray scanner (Nirmidas Biotech, Inc.) was used to scan each biochip for IRDye800-conjugated anti-human IgG and CF-647-conjugated anti-human IgM signals on the SARS-CoV-2 antigen spots. CF-647 and IRDye800 fluorescence images in the respective red and green channels were generated and the median fluorescence signal (MFI) for each microarray was quantified by the MidaScan Software version 2.0.0. The data was used to calculate the average MFI with the antigen spots of the highest and lowest MFIs removed for each channel, thus lending to a single signal intensity used to measure antibody detection in each sample. Afterwards, the MFI was normalized to the average intrawell MFI signal, resulting in an intrawell ratio, and adjusted by a factor of 100 for IgM and 10 for IgG. The final values were used to determine antibody status of the samples for the corresponding antigen. Cutoffs were determined by ROC curve analysis using MedCalc Statistical Software version 19.2.1 (MedCalc Software Ltd., Ostend, Belgium). This method resulted in the optimal combination of sensitivity and specificity of the multiplexed assay on pGOLD.

## Acknowledgements

We thank Lauren Drake of the Loma Linda Medical Center for providing some of the samples used in this work, and Dr. Wenhui Li and Jianhua Sui for providing purified humanized anti-SARS-CoV-2 IgG and for helpful discussions.

## Author Contributions

H.D. and M.T. envisioned and designed the project and experiments. T.L., J.H., S.Z., J.K. and D.S. performed the assay experiments. K.O., D.K., J.R., C. H., C.P., J.B., J.P.R-S., and J.G.M. collected and processed samples. Coauthors from PAMF worked on sample collections and accessing patient information through an IRB. H.D., J.H. and T.L. analyzed the data and wrote the manuscript. All authors commented on the manuscript.

## Additional information

Supplementary information is available for this paper

Correspondence and requests for materials should be addressed to H.D., M.T., or J.G.M.

