## Supplemental Figure 1, 2 and Table 1-8 for "High-Accuracy Multiplexed SARS-CoV-2 Antibody Assay with Avidity and Saliva Capability on a Nano-Plasmonic Platform"

* These authors contributed equally.

**SUPPLEMENTARY INFORMATION**

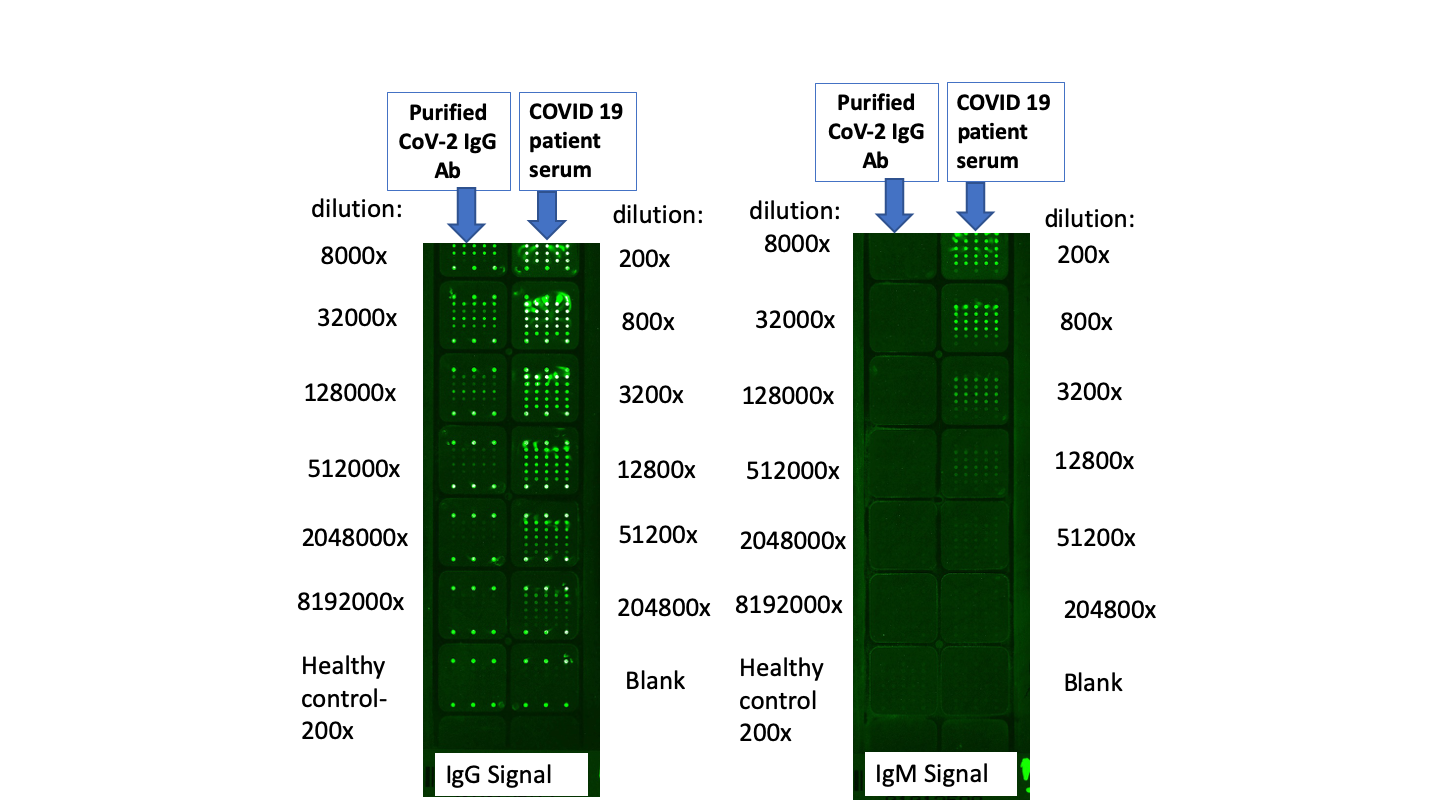

**b**

**a**

**Figure S1.** **Analytical sensitivity of pGOLD SARS-CoV-2 antibody detection.** a) A NIR fluorescence image of a pGOLD biochip (7.5 cm x 2.5 cm) showing SARS-CoV-2 IgG signals labeled by anti-human IgG-IRDye800 of a serially-diluted, purified IgG antibody solution (left column) and a serially-diluted, PCR-confirmed COVID-19 patient serum (right column). b) A NIR fluorescence image of SARS-CoV-2 IgM signals labeled by anti-human IgM-IRDye800 of the serially-diluted, purified IgG antibody (left column) and PCR-confirmed COVID-19 patient serum sample (right column). Each sample dilution was assayed in identical square wells with SARS-CoV-2 antigens S1 and RBD prepared at 5X and 10X and printed in 4 rows with 5 spots each. The top and bottom rows in each well were printed with human IgG as control spots.

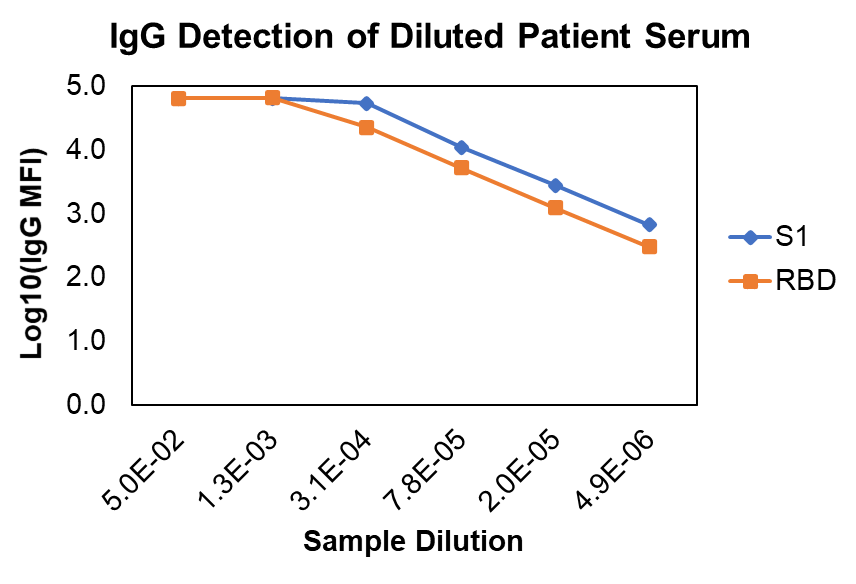

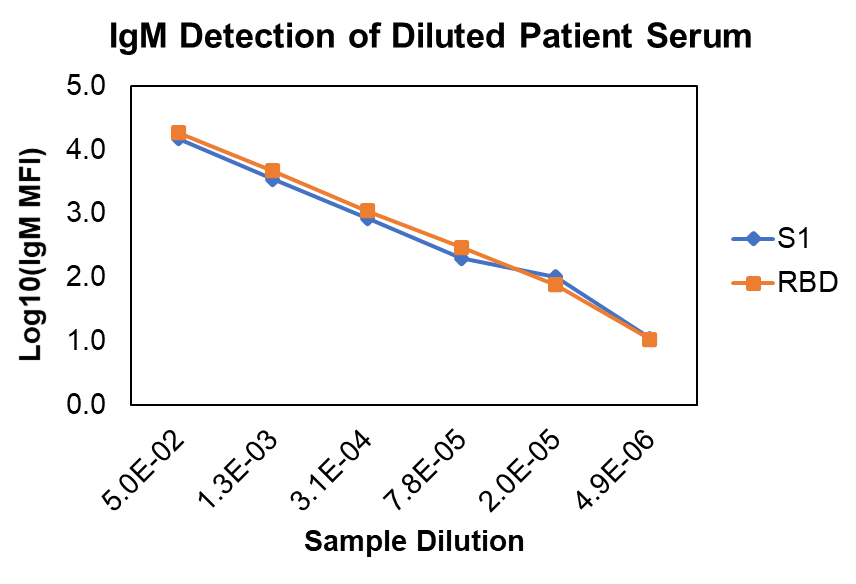

**Figure S2.** Log_10_ of the IgG median fluorescence intensity (MFI) readings (top) and IgM MFI readings (bottom) of a serially-diluted, PCR-confirmed COVID-19 patient serum against SARS-CoV-2 S1 and RBD antigens. A clear IgG signal was seen up to a dilution of 4.9e-06 and a clear IgM signal up to a dilution of 7.8e-05. The IgG signal was saturated at dilutions of 5.0e-02 and 1.3e-03.

| **Number of Samples** | **Origin** | **Sample description** | **pGOLD IgG Results for negative and presumptive negative samples** | **pGOLD IgM Results for negative and presumptive negative samples** |
| --- | --- | --- | --- | --- |
|  |  |  | IgG NEG | IgM NEG |
| 33 | 4 from CDPH and 29 from Loma Linda Medical Center | Confirmed PCR-negative for COVID-19 | 33 | 33 |
| 40 | From Arizona State University Health Services | Pre-pandemic. Samples used as healthy control for projects prior to COVID-19 outbreak | 40 | 40 |
| 311 | JSRLSD at the Palo Alto Medical Foundation | Pre-pandemic  (collected between 2017 to 2019) | 310 | 310 |
| 70 | Pre-pandemic samples for cross-reactivity evaluation | Pre-pandemic samples for cross-reactivity evaluation | 70 | 70 |
| **Specificity** | Total |  | 453/454 = **99.78%** | 453/454= **99.78%** |
| **95% CI:** |  |  | **98.76% - 99.96%** | **98.76% - 99.96%** |

**Table S1.** Negative agreement of pGOLD SARS-CoV-2 IgG/IgM assay against the S1 antigen for specificity using a total of 454 PCR-negative and pre-pandemic cross-reactive or healthy control (“presumptive negative”) samples.

| **Days since symptom onset** | **Number of Samples** | **# IgG Positive** | **# IgM Positive** | **# Antibody Positive** | **% IgG Positive** | **% IgM Positive** | **% Antibody Positive** |
| --- | --- | --- | --- | --- | --- | --- | --- |
| 0 to 7 | 16 | 2 | 7 | 7 | 12.50% | 43.75% | 43.75% |
| 8 to 14 | 21 | 10 | 14 | 14 | 47.62% | 66.67% | 66.67% |
| 15 to 45 | 33 | 33 | 33 | 33 | 100.00% | 100.00% | 100.00% |
| **Total** | 70 | 45 | 54 | 54 |  |  |  |

**Table S2.** Summary of pGOLD SARS-CoV-2 IgG/IgM assay results against the S1 antigen for 70 samples collected from PCR-positive COVID-19 patients at 0 to 7 (Group I), 8 to 14 (Group II), and 15 to 45 (Group III) days since symptom onset.

| **Days since symptom onset** | **Number of PCR+ samples** | **pGOLD assay sensitivity for IgG+** | **pGOLD assay sensitivity for IgM+** | **pGOLD assay sensitivity for IgG+ or IgM+ or both** |
| --- | --- | --- | --- | --- |
| **≥ 6 days** | 60 | 45/60 = **75 %**  (95% CI 62.77%-84.22 %) | 52/60 = **86.67 %**  (95% CI 75.83 % - 93.09%) | 52/60 = **86.67 %**  (95% CI 75.83 % - 93.09%) |
| **≥ 10 days** | 49 | 43/49 = **87.76 %**  (95% CI 75.76%-94.27% ) | 46/49 = **93.87 %**  (95% CI 83.48%-97.89%) | 46/49 = **93.87 %**  (95% CI 83.48%-97.89%) |
| > **14 days** | 33 | 33/33 = **100 %**  (95% CI 89.5%-100%) | 33/33 = **100 %**  (95% CI 89.5%-100%) | 33/33 = **100 %**  (95% CI 89.5%-100%) |

**Table S3.** Positive agreement of pGOLD SARS-CoV-2 IgG/IgM assay against the S1 antigen according to days since PCR-positive COVID-19 serum samples were collected post symptom onset in the range of 6-45 days.

| **Disease Type** | **Reference Method** | **Numbers of Samples** | **Nirmidas pGOLD SARS-CoV-2 IgG/IgM assay (S1)** | | | |
| --- | --- | --- | --- | --- | --- | --- |
|  |  |  | IgM POS | IgM NEG | IgG POS | IgM NEG |
| Coronavirus 229E | BioFire PCR | 1 | 0 | 1 | 0 | 1 |
| Coronavirus NL63 | BioFire PCR | 1 | 0 | 1 | 0 | 1 |
| Coronavirus OC43 | BioFire PCR | 1 | 0 | 1 | 0 | 1 |
| FluA/FluB | ELISA | 5 | 0 | 5 | 0 | 5 |
| FluA | ELISA | 3 | 0 | 3 | 0 | 3 |
| ANA | BioPlex 2200 antibody test | 13 | 0 | 13 | 0 | 13 |
| HBV (AcHBs) | Diasorin Liaison XL | 5 | 0 | 5 | 0 | 5 |
| Haemophilus Influenzae | BioFire PCR | 1 | 0 | 1 | 0 | 1 |
| Rheumatoid Factor | ELISA | 1 | 0 | 1 | 0 | 1 |
| RSV | PCR | 8 | 0 | 8 | 0 | 8 |
| Zika and Dengue IgG | Euroimmune IgG | 8 | 0 | 8 | 0 | 8 |
| Zika IgG/IgM | Euroimmune IgG/InBios IgM | 5 | 0 | 5 | 0 | 5 |
| CHIKV IgG | Euroimmune IgG | 2 | 0 | 2 | 0 | 2 |
| HCV | Antibody test | 6 | 0 | 6 | 0 | 6 |
| HIV | Antibody test | 10 | 0 | 10 | 0 | 10 |
| **Total** |  | **70** | **0** | **70** | **0** | **70** |

**Table S4.**  Cross-reactivity evaluation study conducted for the pGOLD SARS-CoV-2 IgG/IgM assay on the S1 antigen, testing 70 serum samples collected from patients with various diseases. None of the samples tested were positive for either IgG or IgM.

| **Sample source** | **Sample ID** | **RT-PCR Result** | **Days since symptom onset** | **RBD IgM Level** | **RBD IgG Level** | **S1 IgM Level** | **S1 IgG Level** | **RBD IgM Status** | **RBD IgG Status** | **S1 IgM Status** | **S1 IgG Status** |
| --- | --- | --- | --- | --- | --- | --- | --- | --- | --- | --- | --- |
| PAMF | 8 | Positive | 0 | -0.38 | 0.00 | -0.57 | 0.05 | neg | neg | neg | neg |
| PAMF | 5 | Positive | 1 | -0.40 | 0.00 | -0.51 | 0.02 | neg | neg | neg | neg |
| PAMF | 20 | Positive | 1 | 0.02 | 0.00 | -0.18 | 0.06 | neg | neg | neg | neg |
| CDPH | 8 | Positive | 2 | 0.69 | 0.00 | 0.26 | 0.02 | neg | neg | neg | neg |
| PAMF | 12 | Positive | 2 | 8.70 | 1.01 | 4.65 | 0.62 | **pos** | neg | **pos** | neg |
| PAMF | 17 | Positive | 3 | 11.86 | 2.08 | 7.86 | 1.50 | **pos** | **pos** | **pos** | **neg** |
| PAMF | 4 | Positive | 4 | -0.04 | 0.06 | 0.03 | 0.05 | neg | neg | neg | neg |
| PAMF | 21 | Positive | 4 | 0.33 | 0.22 | 0.12 | 0.11 | neg | neg | neg | neg |
| PAMF | 6 | Positive | 5 | -0.33 | 0.00 | -0.32 | 0.01 | neg | neg | neg | neg |
| PAMF | 7 | Positive | 5 | -0.42 | 0.02 | -0.53 | 0.07 | neg | neg | neg | neg |
| CDPH | 2.34 | Positive | 6 | 4.77 | 0.27 | 2.60 | 0.14 | **pos** | neg | **pos** | neg |
| PAMF | 65 | Positive | 6 | -0.37 | 0.03 | 2.00 | 15.67 | **neg** | **neg** | **pos** | **pos** |
| CDPH | 2.37 | Positive | 7 | 72.16 | 28.76 | 36.23 | 11.11 | **pos** | **pos** | **pos** | **pos** |
| PAMF | 15 | Positive | 7 | 9.67 | 1.81 | 7.01 | 1.07 | **pos** | **pos** | **pos** | **neg** |
| PAMF | 74 | Positive | 7 | 5.08 | 1.53 | 3.30 | 1.00 | **pos** | neg | **pos** | neg |
| PAMF | 100 | Positive | 7 | 0.52 | 0.02 | 0.54 | 0.01 | neg | neg | neg | neg |
| CDPH | 16 | Positive | 8 | 0.26 | 0.08 | 0.92 | 0.04 | neg | neg | neg | neg |
| PAMF | 64 | Positive | 8 | 0.02 | 0.07 | -0.45 | 0.06 | neg | neg | neg | neg |
| PAMF | 102 | Positive | 8 | 4.22 | 1.57 | 2.00 | 0.99 | **pos** | neg | **pos** | neg |
| CDPH | 6 | Positive | 9 | 1.34 | 0.50 | 1.37 | 0.37 | neg | neg | neg | neg |
| PAMF | 70 | Positive | 9 | 1.41 | 0.10 | 0.56 | 0.03 | neg | neg | neg | neg |
| CDPH | 2 | Positive | 10 | 29.28 | 7.15 | 24.52 | 6.74 | **pos** | **pos** | **pos** | **pos** |
| CDPH | 2.1 | Positive | 10 | 8.92 | 0.35 | 6.26 | 0.24 | **pos** | neg | **pos** | neg |
| PAMF | 78 | Positive | 10 | 1.17 | 0.12 | 1.24 | 0.18 | neg | neg | neg | neg |
| CDPH | 2.32 | Positive | 11 | 14.56 | 6.02 | 6.97 | 3.64 | **pos** | **pos** | **pos** | **pos** |
| CDPH | 2.18 | Positive | 12 | 421.35 | 41.51 | 421.35 | 41.50 | **pos** | **pos** | **pos** | **pos** |
| CDPH | 2.28 | Positive | 12 | 30.08 | 20.28 | 15.92 | 13.67 | **pos** | **pos** | **pos** | **pos** |
| CDPH | 2.29 | Positive | 12 | 44.93 | 39.90 | 25.35 | 36.20 | **pos** | **pos** | **pos** | **pos** |
| PAMF | 14 | Positive | 12 | 3.58 | 1.08 | 2.75 | 1.44 | **pos** | neg | **pos** | neg |
| PAMF | 73 | Positive | 12 | 3.34 | 0.78 | 2.77 | 1.37 | **pos** | neg | **pos** | neg |
| CDPH | 2.4 | Positive | 13 | 6.86 | 1.22 | 3.10 | 1.67 | **pos** | **neg** | **pos** | **pos** |
| CDPH | 5 | Positive | 13 | 16.53 | 7.03 | 17.72 | 10.82 | **pos** | **pos** | **pos** | **pos** |
| PAMF | 101 | Positive | 13 | 6.61 | 10.27 | 5.15 | 10.33 | **pos** | **pos** | **pos** | **pos** |
| PAMF | 115 | Positive | 13 | -0.08 | 0.01 | -0.02 | -0.03 | neg | neg | neg | neg |
| CDPH | 14 | Positive | 14 | 369.91 | 37.07 | 369.74 | 37.03 | **pos** | **pos** | **pos** | **pos** |
| CDPH | 15 | Positive | 14 | 17.41 | 16.06 | 15.74 | 19.76 | **pos** | **pos** | **pos** | **pos** |
| PAMF | 114 | Positive | 14 | 1.07 | 0.14 | 1.13 | 0.21 | neg | neg | neg | neg |
| CDPH | 2.6 | Positive | 15 | 94.42 | 39.11 | 75.64 | 33.74 | **pos** | **pos** | **pos** | **pos** |
| CDPH | 7 | Positive | 15 | 63.92 | 37.08 | 64.09 | 37.01 | **pos** | **pos** | **pos** | **pos** |
| CDPH | 9 | Positive | 15 | 349.33 | 35.09 | 349.14 | 35.01 | **pos** | **pos** | **pos** | **pos** |
| CDPH | 2.8 | Positive | 16 | 88.08 | 41.25 | 75.82 | 41.21 | **pos** | **pos** | **pos** | **pos** |
| CDPH | 2.21 | Positive | 17 | 274.49 | 43.33 | 156.57 | 43.24 | **pos** | **pos** | **pos** | **pos** |
| CDPH | 2.22 | Positive | 17 | 252.71 | 41.68 | 137.63 | 41.63 | **pos** | **pos** | **pos** | **pos** |
| CDPH | 2.27 | Positive | 17 | 48.47 | 36.60 | 26.60 | 36.56 | **pos** | **pos** | **pos** | **pos** |
| CDPH | 2.31 | Positive | 18 | 48.64 | 45.90 | 35.45 | 45.85 | **pos** | **pos** | **pos** | **pos** |
| CDPH | 2.36 | Positive | 18 | 249.57 | 46.82 | 157.49 | 46.70 | **pos** | **pos** | **pos** | **pos** |
| PAMF | 13 | Positive | 18 | 16.84 | 21.36 | 17.48 | 22.86 | **pos** | **pos** | **pos** | **pos** |
| CDPH | 2.26 | Positive | 19 | 99.78 | 36.23 | 59.71 | 36.18 | **pos** | **pos** | **pos** | **pos** |
| PAMF | 10 | Positive | 19 | 36.68 | 39.18 | 31.64 | 39.12 | **pos** | **pos** | **pos** | **pos** |
| PAMF | 16 | Positive | 19 | 48.35 | 34.91 | 43.81 | 34.81 | **pos** | **pos** | **pos** | **pos** |
| CDPH | 2.13 | Positive | 20 | 219.25 | 37.47 | 235.55 | 37.42 | **pos** | **pos** | **pos** | **pos** |
| CDPH | 2.24 | Positive | 20 | 147.42 | 38.29 | 84.09 | 38.25 | **pos** | **pos** | **pos** | **pos** |
| CDPH | 2.3 | Positive | 20 | 126.42 | 41.16 | 63.92 | 41.09 | **pos** | **pos** | **pos** | **pos** |
| CDPH | 2.25 | Positive | 21 | 124.12 | 33.79 | 57.16 | 33.72 | **pos** | **pos** | **pos** | **pos** |
| CDPH | 2.2 | Positive | 22 | 69.31 | 36.77 | 68.65 | 36.69 | **pos** | **pos** | **pos** | **pos** |
| CDPH | 2.35 | Positive | 22 | 37.40 | 43.17 | 53.51 | 43.11 | **pos** | **pos** | **pos** | **pos** |
| CDPH | 2.19 | Positive | 23 | 91.08 | 37.18 | 68.09 | 37.11 | **pos** | **pos** | **pos** | **pos** |
| PAMF | 9 | Positive | 23 | 80.63 | 41.34 | 50.69 | 41.26 | **pos** | **pos** | **pos** | **pos** |
| PAMF | 11 | Positive | 23 | 10.13 | 14.00 | 5.20 | 13.01 | **pos** | **pos** | **pos** | **pos** |
| PAMF | 22 | Positive | 23 | 13.28 | 13.26 | 6.38 | 12.05 | **pos** | **pos** | **pos** | **pos** |
| CDPH | 2.38 | Positive | 25 | 69.01 | 42.79 | 44.57 | 42.68 | **pos** | **pos** | **pos** | **pos** |
| CDPH | 2.12 | Positive | 26 | 224.75 | 36.07 | 55.89 | 35.21 | **pos** | **pos** | **pos** | **pos** |
| CDPH | 2.7 | Positive | 26 | 59.84 | 38.48 | 56.78 | 38.43 | **pos** | **pos** | **pos** | **pos** |
| CDPH | 2.14 | Positive | 27 | 20.75 | 39.08 | 21.91 | 39.01 | **pos** | **pos** | **pos** | **pos** |
| CDPH | 2.16 | Positive | 27 | 195.37 | 40.50 | 79.72 | 40.39 | **pos** | **pos** | **pos** | **pos** |
| CDPH | 2.17 | Positive | 27 | 133.82 | 37.44 | 76.70 | 37.36 | **pos** | **pos** | **pos** | **pos** |
| CDPH | 2.11 | Positive | 32 | 47.91 | 41.14 | 44.62 | 41.09 | **pos** | **pos** | **pos** | **pos** |
| CDPH | 2.15 | Positive | 36 | 42.79 | 38.77 | 34.18 | 38.72 | **pos** | **pos** | **pos** | **pos** |
| PAMF | 18 | Positive | 38 | 157.27 | 36.24 | 90.97 | 36.20 | **pos** | **pos** | **pos** | **pos** |
| CDPH | 2.9 | Positive | 45 | 29.91 | 42.48 | 31.74 | 42.43 | **pos** | **pos** | **pos** | **pos** |

**Table S5.** Comparison of pGOLD SARS-CoV-2 IgG/IgM assay results of PCR-positive patient samples (with known data on days since symptom onset) against the S1 and RBD antigens. The IgG positive status of a sample was determined if the IgG level was > 1.79 and > 1.62 for RBD and S1, respectively. The IgM positive status of a sample was determined if the IgM level was > 3 and > 1.38 for RBD and S1, respectively.

| **Days since symptom onset** | **Number of samples** | **# IgG Positive** | **# IgM Positive** | **# Antibody Positive** | **% IgG Positive** | **% IgM Positive** | **% Antibody Positive** |
| --- | --- | --- | --- | --- | --- | --- | --- |
| 0 to 7 | 16 | 3 | 6 | 6 | 18.75% | 37.50% | 37.50% |
| 8 to 14 | 21 | 9 | 14 | 14 | 42.86% | 66.67% | 66.67% |
| 15 to 45 | 33 | 33 | 33 | 33 | 100.00% | 100.00% | 100.00% |
| **Total** | 70 | 45 | 53 | 53 |  |  |  |

**Table S6.** Summary of pGOLD SARS-CoV-2 IgG/IgM assay results against the RBD antigen for 70 samples collected from PCR-positive COVID-19 patients at 0 to 7 (Group I), 8 to 14 (Group II), and 15 to 45 (Group III) days since symptom onset.

| **Days from symptom onset date** | **Number of PCR+ samples** | **pGOLD assay sensitivity for IgG+** | **pGOLD assay sensitivity for IgM+** | **pGOLD assay sensitivity for IgG+ or IgM+ or both** |
| --- | --- | --- | --- | --- |
| **≥ 6 days** | 60 | 44/60 = **73.33 %**  (95% CI 57.64%-79.76%) | 51/60 = **85 %**  (95% CI 73.89% -91.90%) | 51/60 = **85 %**  (95% CI 73.89% -91.90%) |
| **≥ 10 days** | 49 | 42/49 = **85.71 %**  (95% CI 73.33%-92.90%) | 46/49 = **93.87 %**  (95% CI 83.48%-97.89%) | 46/49 = **93.87 %**  (95% CI 83.48%-97.89%) |
| > **14 days** | 33 | 33/33 = **100 %**  (95% CI 89.5%-100%) | 33/33 = **100 %**  (95% CI 89.5%-100%) | 33/33 = **100 %**  (95% CI 89.5%-100%) |

**Table S7.** Positive agreement of pGOLD SARS-CoV-2 IgG/IgM assay against the RBD antigen according to days since PCR-positive COVID-19 serum samples were collected post symptom onset in the range of 6-45 days.

| **Sample source** | **Sample ID** | **Days since symptom onset** | **S1 Untreated IgG MFI** | **S1 6M Urea-treated IgG MFI** | **S1 Avidity Index** | **RBD Untreated IgG MFI** | **RBD 6M Urea-treated IgG MFI** | **RBD Avidity Index** |
| --- | --- | --- | --- | --- | --- | --- | --- | --- |
| PAMF | 65 | 6 | 10456.17 | 8872.33 | 0.85 | NA | NA | NA |
| CDPH | 2.37 | 7 | 707.17 | 10.00 | 0.01 | 2214.50 | 4.33 | 0.00 |
| PAMF | 15 | 7 | 512.83 | 14.00 | 0.03 | 743.00 | 4.67 | 0.01 |
| PAMF | 102 | 8 | 515.50 | 17.00 | 0.03 | 514.17 | 8.33 | 0.02 |
| CDPH | 2 | 10 | 1675.17 | 19.33 | 0.01 | 1182.33 | -36.83 | -0.03 |
| CDPH | 2.32 | 11 | 870.67 | 14.33 | 0.02 | 1220.83 | 12.33 | 0.01 |
| CDPH | 2.29 | 12 | 19303.50 | 379.33 | 0.02 | 216.83 | 4.67 | 0.02 |
| PAMF | 14 | 12 | 324.67 | 17.33 | 0.05 | 5987.00 | 3.67 | 0.00 |
| CDPH | 2.28 | 12 | 5193.83 | 25.67 | 0.00 | 217.50 | 1.17 | 0.01 |
| PAMF | 73 | 12 | 410.67 | 18.00 | 0.04 | 19612.17 | 94.33 | 0.00 |
| CDPH | 2.18 | 12 | 45554.00 | 7471.50 | 0.16 | 39149.17 | 204.33 | 0.01 |
| CDPH | 5 | 13 | 3136.00 | 26.67 | 0.01 | 319.33 | 2.00 | 0.01 |
| CDPH | 2.4 | 13 | 621.67 | 10.67 | 0.02 | 2091.17 | 4.33 | 0.00 |
| PAMF | 101 | 13 | 4829.17 | 118.67 | 0.02 | 2550.00 | 27.33 | 0.01 |
| CDPH | 15 | 14 | 5017.83 | 319.00 | 0.06 | 24448.50 | 33.00 | 0.00 |
| CDPH | 14 | 14 | 36119.83 | 3098.67 | 0.09 | 3833.33 | 6.00 | 0.00 |
| CDPH | 7 | 15 | 35329.17 | 1376.00 | 0.04 | 37507.17 | 104.00 | 0.00 |
| CDPH | 2.6 | 15 | 7136.83 | 153.83 | 0.02 | 33893.83 | 32.00 | 0.00 |
| CDPH | 9 | 15 | 44496.00 | 3858.33 | 0.09 | 5862.67 | -30.33 | -0.01 |
| CDPH | 2.8 | 16 | 53572.17 | 3365.17 | 0.06 | 54150.33 | 99.00 | 0.00 |
| CDPH | 2.21 | 17 | 35587.83 | 10853.83 | 0.30 | 16938.50 | 263.83 | 0.02 |
| CDPH | 2.22 | 17 | 65442.33 | 6419.17 | 0.10 | 57113.17 | 299.67 | 0.01 |
| CDPH | 2.27 | 17 | 65431.67 | 2070.67 | 0.03 | 65437.33 | 98.33 | 0.00 |
| PAMF | 13 | 18 | 4680.00 | 318.50 | 0.07 | 65450.00 | 3662.83 | 0.06 |
| CDPH | 2.31 | 18 | 65446.67 | 4374.00 | 0.07 | 58531.67 | 6077.33 | 0.10 |
| CDPH | 2.36 | 18 | 60468.00 | 5845.17 | 0.10 | 4650.33 | 221.00 | 0.05 |
| PAMF | 16 | 19 | 64600.33 | 4361.33 | 0.07 | 47921.00 | 4972.33 | 0.10 |
| CDPH | 2.26 | 19 | 44727.00 | 1193.67 | 0.03 | 40300.50 | 4376.00 | 0.11 |
| PAMF | 10 | 19 | 46299.17 | 4255.83 | 0.09 | 34217.67 | 100.00 | 0.00 |
| CDPH | 2.3 | 20 | 50803.83 | 2773.50 | 0.05 | 36666.50 | 4893.83 | 0.13 |
| CDPH | 2.13 | 20 | 65437.67 | 6894.00 | 0.11 | 38427.50 | 193.17 | 0.01 |
| CDPH | 2.24 | 20 | 51132.00 | 1832.17 | 0.04 | 55682.67 | 261.33 | 0.00 |
| CDPH | 2.25 | 21 | 65433.33 | 5994.83 | 0.09 | 58754.00 | 297.00 | 0.01 |
| CDPH | 2.35 | 22 | 65431.67 | 9695.33 | 0.15 | 65438.00 | 10464.17 | 0.16 |
| CDPH | 2.2 | 22 | 65335.00 | 4239.33 | 0.06 | 64295.83 | 232.33 | 0.00 |
| PAMF | 11 | 23 | 2868.00 | 418.67 | 0.15 | 46869.33 | 3470.67 | 0.07 |
| PAMF | 9 | 23 | 31018.33 | 3122.33 | 0.10 | 6044.00 | 346.17 | 0.06 |
| PAMF | 22 | 23 | 4733.67 | 217.33 | 0.05 | 3396.00 | 563.50 | 0.17 |
| CDPH | 2.19 | 23 | 43545.83 | 2331.00 | 0.05 | 26088.83 | 104.50 | 0.00 |
| CDPH | 2.38 | 25 | 22467.67 | 692.00 | 0.03 | 25060.67 | 852.00 | 0.03 |
| CDPH | 2.7 | 26 | 64538.33 | 5847.17 | 0.09 | 54204.00 | 290.00 | 0.01 |
| CDPH | 2.12 | 26 | 7071.33 | 87.00 | 0.01 | 7509.83 | 37.33 | 0.00 |
| CDPH | 2.14 | 27 | 17324.00 | 2569.17 | 0.15 | 8868.00 | 99.33 | 0.01 |
| CDPH | 2.16 | 27 | 65426.67 | 5988.33 | 0.09 | 16948.33 | 106.67 | 0.01 |
| CDPH | 2.17 | 27 | 24028.50 | 1617.67 | 0.07 | 62232.33 | 298.50 | 0.00 |
| CDPH | 2.11 | 32 | 63233.67 | 8605.00 | 0.14 | 46867.83 | 229.67 | 0.00 |
| CDPH | 2.15 | 36 | 33753.33 | 2229.50 | 0.07 | 22320.33 | 101.00 | 0.00 |
| PAMF | 18 | 38 | 27269.50 | 849.00 | 0.03 | 56669.67 | 550.50 | 0.01 |
| CDPH | 2.9 | 45 | 59050.33 | 12739.33 | 0.22 | 38435.17 | 405.83 | 0.01 |

**Table S8.** pGOLD SARS-CoV-2 IgG avidity assay results of IgG-positive, PCR-confirmed COVID-19 patient samples (in the range of 6-45 days post symptom onset) against the S1 and RBD antigens. The samples were treated with and without 6M urea and an avidity index was calculated by dividing the IgG signals of the urea-treated samples by the IgG signals of the non-treated sample for the respective antigen.
